# A Dimensionality Reduction Approach for Motor Imagery BCI using Functional Clustering, Graph Signal Processing, and Differential Evolution

**DOI:** 10.1101/2023.08.28.555094

**Authors:** Mohammad Davood Khalili, Vahid Abootalebi, Hamid Saeedi-Sourck

## Abstract

This paper aims to address the dimension reduction and classification of electroencephalogram (EEG) signals within the context of motor imagery brain-computer interface (MI-BCI). By leveraging modern brain signal processing tools, specifically Graph Signal Processing (GSP) and meta-heuristic techniques, we introduce the K-GLR-DE approach. This methodology encompasses functional clustering, Kron reduction, regularized common spatial patterns with generic learning (GLRCSP), and differential evolution (DE). Our approach is underpinned by a comprehensive structural-functional framework that carefully shapes the architecture of the brain graph. Edge weights are assigned based on geometric distance and correlation, imbuing the model with physiologically meaningful connectivity patterns. Graph reduction involves strategically employing physiological regions of interest (ROIs) and Kron reduction to select informative subgraphs while preserving vital information from all graph vertices. Feature extraction integrates total variation calculation and the GLRCSP method, followed by dimension reduction using the DE algorithm. The extracted features are then evaluated using well-established machine-learning classifiers. The validation process is carried out using Dataset IVa from BCI Competition III, providing a tangible benchmark for the performance of the K-GLR-DE approach. Significantly, the SVM-RBF classifier stands out as the top performer, achieving a remarkable average accuracy of 96.46±0.83. Noteworthy is our approach’s capacity to notably augment MI-BCI classification performance across diverse training trial scenarios, encompassing limited, small, and conventional settings.

## 1. Introduction

A Brain-Computer Interface (BCI) serves as a communication system in which user communication bypasses traditional output pathways of the brain, specifically peripheral nerves and muscles [1]. Fundamentally, a BCI empowers subjects to communicate and control using their brain signals [2-4]. Electroencephalogram (EEG) is the preferred choice for BCI applications due to its cost-effectiveness, high temporal resolution, non-invasiveness, and user-friendly nature in comparison to alternative methods [5]. EEG-based non-invasive BCI systems are typically organized into five primary categories: slow cortical potentials, motor imagery (MI), P300 event-related potential (P300 ERP), steady-state visual evoked potential, and error-related negative evoked potential [6]. Most BCI systems necessitate signals from multiple locations across the scalp to optimize performance [7]. However, an excessive number of EEG channels may introduce redundancy and potentially undermine BCI performance [8, 9]. Thus, the strategic selection of an adequate number of channels that balance accuracy and practicality becomes crucial [7]. In MI-BCI, subjects mentally simulate precise motor actions without physically executing the movements. Subsequently, the BCI system discerns the neural activity associated with this mental simulation and encodes the EEG signal. This process helps subjects acquire the neural patterns associated with specific movements, enabling them to learn the corresponding physical action more effectively.

Advances in data science have increased the interest in complex signals with irregular structures [10]. High-dimensional data can live on weighted graph vertices and show the geometric structure of data in various applications, such as social networks, energy systems, transportation, sensors, and neural networks [11]. Graph Signal Processing (GSP) is an innovative tool for analyzing data on complex and irregular structures, with particular applications in brain signals [12]. GSP enhances signal processing through a graph-based perspective integrated with classical signal processing, providing enhanced insight into signal connectivity effects [13]. GSP encompasses methods such as graph Fourier transform (GFT) [12-14], graph shift [12, 13], graph filter [11, 13, 15], and spectral graph wavelet transform (SGWT) [15-17] for processing graph-based signals.

Numerous studies within brain signal analysis have employed conventional machine learning techniques like principal component analysis (PCA), independent component analysis (ICA), and support vector machine (SVM). Recent studies have utilized graph-based frameworks and GSP perspectives, enhancing our understanding of brain phenomena and interconnectivity among brain regions. These investigations have provided insights into brain signal classification [18, 19], diagnosis of brain diseases [20-23], BCI [24-26], and graph frequency analysis of brain signals [27].

In recent years, there has been several research endeavors aimed at reducing the dimensions of BCI-EEG data using GSP techniques. Tanaka et al. [24] pursued the dimensionality reduction of sample covariance matrices (SCM) in BCI applications by formulating a geometric graph based on the distribution of electrodes across the head and investigating structural connections using geographical distance measures. For accurate and low-dimensional SCM estimation in the tangent space mapping (TSM) method, they employed GFT-based dimension reduction. This approach extracted SCM from the reduced data by leveraging the first *r* eigenvectors of the graph Laplacian matrix, effectively reducing the column space [24]. Kalantar et al. [25] constructed a geometric graph to examine the structural and functional connections in the brain. They derived edge weights for the BCI application by combining geographical distance and Pearson correlation among EEG electrodes. They employed GFT, TSM, and PCA for dimension reduction, feature extraction, and feature selection, respectively [25]. In another study, Kalantar et al. [26] developed a hybrid graph for BCI applications. They performed graph clustering and selected the top two clusters using linear discriminant analysis (LDA). They extracted features using common spatial patterns (CSP) and then inputted these features into LDA, quadratic discriminant analysis (QDA), and logistic regression (LR) classifiers. We constructed structural-functional graphs similar to [25], albeit with different normalization and optimization coefficients [28]. The dimensionality of these graphs was efficiently reduced through a fusion of Kron reduction and GFT. Statistical feature extraction was conducted using the Ledoit-Wolf shrinkage estimator and TSM on EEG data from superior vertices. The dimension reduction of these features was achieved by PCA and an optimization technique known as differential evolution (DE).

Prior studies [24-26, 28] have revealed various challenges that hinder the improvement of BCI application outcomes. The primary contributions of this paper include:

1. Efficiently reducing the dimensions of extensive EEG data within a complex brain graph framework.
2. Thoughtfully selecting an excellent channel subset for high performance.
3. Employing a processing methodology that ensures high classification accuracy and efficiency for online BCI applications, along with an appropriate test execution speed.
4. Developing a processing approach suitable for training datasets of varying sizes, including limited, small, and conventional datasets, with adjustable parameters for improved decoding accuracy.

This article addresses the aforementioned contributions and analyzes the corresponding results. To address the first and second contributions, we suggest employing a combination of Kron reduction and functional clustering. This approach effectively reduces the dimensions of BCI data by leveraging the perspective of GSP. Furthermore, we utilize the DE algorithm to identify and select superior features. To address the third and fourth contributions, we propose an approach that incorporates Kron reduction, regularized common spatial patterns with generic learning (GLRCSP) for feature extraction, and DE for feature selection. Throughout different stages, we optimize and adjust parameters for each subject.

The paper is organized as follows: Section 2 presents an overview of the dataset, introduces fundamental concepts for GSP and brain graph, and provides preliminaries on Kron reduction, RCSP, and differential evolution. Section 3 outlines the comprehensive process of our proposed method, encompassing preprocessing, feature extraction, dimension reduction, feature selection, and classification steps. Section 4 presents the simulation results and subsequent discussions, while Section 5 serves as the conclusion of the paper.

## 2. Materials and Methods

### 2.1. Dataset

The Intelligent Data Analysis Group [29] provided Dataset IVa from BCI competition III. This publicly available dataset includes EEG signals recorded during Motor Imagery tasks involving the right hand and right foot. EEG data were collected from five healthy subjects using 118 electrodes based on the extended international 10-20 system. The signals were filtered within the range of (0.05-200 Hz) and digitized at a sampling rate of 1000 Hz. The dataset comprises data from the initial four sessions conducted without any feedback. It encompasses two types of visual stimulation: (1) Target letters presented behind a fixation cross, potentially inducing slight target-correlated eye movements, and (2) Targets displayed as randomly moving objects, potentially causing target-uncorrelated eye movements. Visual cues were presented for 3.5 seconds during each trial. Each subject performed a total of 280 MI trials involving either the right hand (R) or foot (F) MI tasks. The presentation of target cues was randomly interrupted by resting intervals lasting between 1.75 to 2.25 seconds. This strategic interruption prevents the formation of temporal dependencies, thereby ensuring the independence of each trial [29, 30].

This paper incorporates training trials for all subjects, utilizing three distinct modes: limited (60), small (100), and conventional (200). Additionally, a specific training trial mode was formulated to align with the BCI Competition III-Dataset IVa (as presented in Table 1) to ensure unbiased comparability with approaches employed in this competition.

**Table 1:** Train and Test Trials in BCI Competition III-Dataset IVa.

| Subject | aa | al | av | aw | ay |
| --- | --- | --- | --- | --- | --- |
| Train Trials | 168 | 224 | 84 | 56 | 28 |
| Test Trials | 112 | 56 | 196 | 224 | 252 |

### 2.2. Graph Signal Processing

A weighted graph *𝒢* is represented as *𝒢* = (*𝒱, ℰ*, ***W***), where *𝒱, ℰ*, and ***W*** denote vertices, edges, and weighted adjacency matrix, respectively. The set *𝒱* = {1, 2, …, *N*} represents vertices, and *ℰ* is a set of edges with tuples (*i, j*) that reflects the connectivity between the *i*^th^ and *j*^th^ nodes. We focus on connected and undirected graphs without self-loops or multiple edges. The Laplacian and normalized Laplacian matrices are defined as ***L*** = ***D*** − ***W***, and ***L***_*n*_ = ***D***^−1*/*2^ ***LD***^−1*/*2^, respectively. Here, ***W*** is the weighted adjacency matrix, and ***D*** is the diagonal degree matrix with diagonal elements 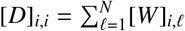. In the case of an undirected graph, ***L*** is a real symmetric positive semi-definite matrix with nonnegative eigenvalues 0 = *λ*_1_ *< λ*_2_ ≤ …≤ *λ*_*N*_, and the corresponding real-valued orthonormal eigenvectors 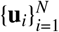. Therefore, ***L*** can be decomposed as ***L*** = ***U***Λ***U***^*T*^, where Λ = diag{*λ*_*l*_} and the columns of ***U*** are 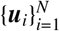. In this context, ***x*** = [*x*(1), …, *x*(*N*)] represents a graph signal, where sample *x*(*m*) is associated with the *m*th node. Furthermore, GFT of a given signal ***x*** over an undirected graph is defined as 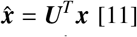 [11]. Equivalently, the inverse GFT is represented as 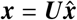.

Total variation (TV) is a widely used graph measure that quantifies signal smoothness concerning the graph. *TV*_𝒢_(***x***) represents the extent of signal changes within the graph, where smaller values indicate slower changes [12]. The total variation of a graph signal ***x*** is defined as [31]:

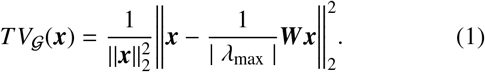

Using (1), we calculate the sum of graph signal variations at each time instant for all time samples in all trials. Here, ***x*** represents the vector of values obtained from all vertices at a specific time, ***W*** denotes the weighted adjacency matrix, and *λ*_max_ corresponds to the largest-magnitude eigenvalue of ***W***.

### 2.3. Brain Graph

Feature extraction, dimension reduction, and classification of brain signals form fundamental aspects in the design of BCI systems. In this study, we employ a graph-based approach to process brain signals, leveraging the rich information provided by the brain graph. The brain graph is defined by vertices and weighted edges, which represent the interconnections among various brain regions. By applying GSP techniques to the signal on the brain graph, we extract relevant features and reduce data dimensionality. GSP-based approaches exhibit remarkable versatility in brain signal processing, as demonstrated by their diverse applications, including: 1) Dimension reduction of EEG signals [24-26, 28], 2) Inference of brain graph topology [10, 27], 3) Spectral analysis of brain signals using wavelet transform [17, 32], 4) Analysis of dynamic brain connectivity [17, 33], and 5) Classification of ERPs [27, 34]. By considering the intricate interplay between brain signals and the underlying graph, GSP enables the integration of physiological and anatomical information. This integration enhances our understanding of brain function and facilitates the development of more robust BCI systems.

The comprehensive analysis of a brain graph requires considering five essential elements: 1) Vertex characteristics, 2) Edge weights that measure the relationship between vertices, 3) Brain graph topology and topological metrics, 4) Brain graph dynamics, and 5) Interconnection relationships across multiple domains and scales [35]. This research focuses on the first two crucial elements of the brain graph: vertex characteristics and edge weights.

Investigating brain graph connectivity requires careful consideration of the composition of brain graph nodes. Brain graph nodes can comprise genes, synapses, neurons, specific brain regions, and brain regions at different spatial scales, reflecting their biological nature and spatial organization. The brain graph incorporates three distinct weighting schemes: structural connectivity, functional connectivity, and effective connectivity. Geometric distance commonly serves as the primary criterion for generating a brain graph based on structural connectivity. Functional connectivity metrics can be categorized into two types: undirected and directed. Undirected measures encompass correlation, coherence, and mutual information, whereas directed metrics encompass Kullback-Leibler divergence, transfer entropy, and phase slope index. Effective connectivity, which captures causal interactions, represents the third aspect of connectivity. The most practical methodologies in this context are Granger causality modeling and dynamic causal modeling, which yield additional insights into the brain graph [35].

Invasive electrophysiological techniques enable the registration of activity from individual cells, including action potentials and spike trains, and activities from groups of cells, such as local field potentials, facilitating connectivity analysis at a small-scale [36]. In contrast, non-invasive techniques such as EEG, functional magnetic resonance imaging (fMRI), magnetoencephalography (MEG), and functional near-infrared spectroscopy (fNIRS) enable connectivity analysis at a large-scale [37]. In this study, EEG signals are assigned to the vertices of the brain graph, contributing to the construction of a comprehensive large-scale brain graph.

### 2.4. Kron Reduction

Graph simplification methods can be broadly categorized into two general groups [38]. The first group consists of graph sparsification approaches that aim to reduce the number of edges in the graph. Well-known examples include spanners, cuts and spectral sparsifiers [39], as well as methods for solving symmetric, diagonally dominant (SDD) linear systems [40]. The second group comprises graph reduction approaches that focus on decreasing the number of graph vertices. Notable methods in this category include graph coarsing [38, 41], and Kron reduction [42, 43].

Kron reduction is a highly notable graph simplification method in the vertex domain that has been applied in recent years for simplifying electrical circuits [44], power systems [42, 44], and graph structures [43]. Given a weighted and undirected graph 𝒢 = (𝒱, *ℰ*, ***W***) with vertices 𝒱, If ***L*** represents the Laplacian matrix, and 𝒱_1_ is a subset containing at least two vertices from 𝒱, the reduced graph by the Kron reduction can be defined as 𝒢^*Kron*^ = {𝒱_1_, *ℰ*^*Kron*^, ***W***^*Kron*^}. Furthermore, based on [43] and Schur complement theorem [45], considering 𝒱_1_ as the selected vertices and its complement, 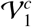 as the deleted vertices, the reduced Laplacian matrix obtained through Kron reduction and the Schur complement of the deleted vertices block from the original Laplacian matrix can be defined as follows:

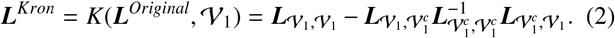

In (2), ***L***_*A,B*_ denotes the submatrix |***A***| × |***B***|, encompassing all elements of matrix ***L*** with row indices in block ***A*** and column indices in block ***B***. Moreover, the first term represents the Laplacian of the selected vertices, while the second term indicates the impact of the Laplacian of the deleted vertices on the Laplacian of the selected vertices. One of the key advantages of Kron reduction is its ability to utilize weights and information from deleted vertices to construct the final Laplacian matrix for the selected vertices. In line with the reduced Laplacian matrix obtained through Kron reduction, the weight of the new graph edges is defined as [42, 43]:

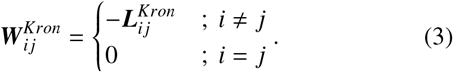

### 2.5. CSP and RCSP

The CSP approach is a widely used method for EEG feature extraction. It discriminates EEG signals or features into two distinct patterns from two classes by simultaneously diagonalizing two real symmetric matrices. One class exhibits the highest variance, while the other class shows the lowest variance [46]. Spatial filters have been shown to effectively improve spatial resolution and increase SNR in EEG signals [46]. The CSP approach, a well-known spatial filtering method, is commonly employed in pattern recognition and BCI systems [47]. By utilizing only a few filters with good separability, the data size is reduced, leading to improved classification accuracy. The objective function *J*_*CS P*_(***W***) is optimized by CSP with spatial filters ***W*** as follows:

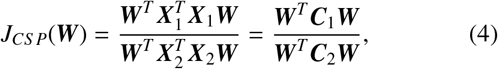

Where *T* denotes transpose, ***X***_1_ and ***X***_2_ correspond to the data matrices of class 1 and class 2 respectively, and ***C***_1_ and ***C***_2_ correspond to the covariance matrices of class 1 and class 2. However, the CSP approach has two disadvantages: sensitivity to noise and susceptibility to overfitting with small training data. The CSP approach requires estimating the spatial covariance matrix for each class, which, in turn, relies on having a clean and sufficient EEG training set; otherwise, the resulting covariance matrices may poorly represent the mental states and lead to ineffective spatial filters [48]. To address these drawbacks, regularized CSP (RCSP) methods have emerged. As proposed in [48], regularizing CSP by incorporating prior information can be useful. As a result, a proper estimation for the covariance matrix is made by integrating prior knowledge and regularization terms as [48]:

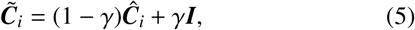

where

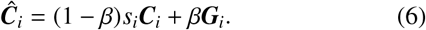

In (5) and (6), ***C***_*i*_ is the initial spatial covariance matrix for class i and 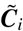 is its regularized estimation. Furthermore, ***I*** represents the identity matrix, and *s*_*i*_ is a scaling parameter. The regularization process involves two parameters, *γ* and *β*, both constrained within the range [0, 1]. Additionally, ***G***_*i*_ represents the generic covariance matrix. The parameter *γ* is employed to shrink the ***C***_*i*_ estimate towards ***I***, effectively mitigating the estimation bias stemming from small training data. Conversely, the parameter *β* facilitates the shrinkage of the ***C***_*i*_ estimate towards ***G***_*i*_, enhancing the stability of the estimation. ***G***_*i*_ signifies the ideal covariance matrix configuration corresponding to each distinct mental state. The acquisition of ***G***_*i*_ involves utilizing signals from diverse subjects who participated same experiments [48]. The application of these methods involves training the spatial filters through the substitution of ***C***_1_ and ***C***_2_ in the conventional CSP algorithm with their regularized counterparts, 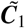 and 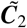. Furthermore, a collection of RCSP approaches is determined based on ***G***_*i*_ and the utilization of one or both adjustment parameters.

GLRCSP leverages covariance matrix regularization using cross-participant data. This approach applies regularization terms *β* and *γ*, which steer the covariance matrix toward both ***I*** and ***G***_*i*_. This dual effect enhances estimation by mitigating bias (*γ*) and bolstering stability (*β*). ***G***_*i*_ is computed through covariance matrices of other subjects, defined as follows:

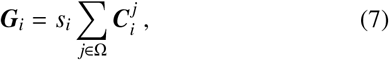

where

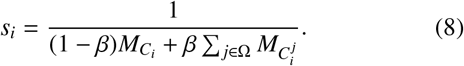

In (7) and (8), *s*_*i*_ is the scaling parameter, Ω denotes the dataset of subjects, 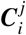 signifies the spatial covariance matrix for class i and subject j, and *M*_*C*_ represents the number of trials utilized to compute the covariance matrix ***C***.

### 2.6. Differential Evolution

Meta-heuristic methods provide a strategic response to addressing the dimensionality challenges by intelligently selecting features based on evaluations of classification performance. These algorithms bear significant promise, having demonstrated success in effectively navigating expansive feature spaces to uncover optimal solutions across diverse applications. Notable among these meta-heuristic techniques are particle swarm optimization [49, 50], differential evolution (DE) [49, 50], artificial bee colony [49], ant colony optimization [49], genetic algorithms (GA) [50], and firefly algorithm [51]. This assortment has proven effective in both feature selection and dimensionality reduction [52].

The DE method emerges as a prominent contender among evolutionary optimization techniques. DE method, characterized by its straightforward yet robust population-based random search algorithm, has demonstrated effective application in addressing optimization challenges spanning diverse fields of basic sciences and engineering. DE offers an array of strategies to generate trial vectors, each with distinct suitability for addressing specific problems. Three pivotal control parameters—population size (n), scale factor (F), and crossover rate (CR)—are integral components of this method, profoundly influencing the effectiveness of the DE approach. As an evolutionary process, the DE algorithm focuses on a population of size n denoted as 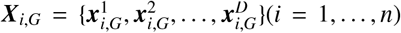 and encodes potential solutions. This population consists of n-dimensional parametric D vectors known as individuals and converge toward the globally optimal direction [53]. Analogous to the GA approach, the DE optimization method employs vector-based search techniques involving crossover and mutation operations. DE boasts two primary strengths: high convergence ability and global optimization capability while overcoming local optimization traps. This study has utilized the most prevalent variant of DE, which encompasses the following procedural steps:

1. Initialization: The algorithm commences by initializing a random population comprising target vectors ***x***_*i*_(*i* = 1, …, *n*).
2. Mutation: For the entire population (*i* = 1, …, *n*), three distinct vectors ***x***_*a*_, ***x***_*b*_, and ***x***_*c*_ are selected randomly. According to (9), a new donor vector is generated by applying a mutation to the *i*th component of the donor vector.

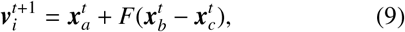

where 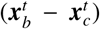 represents the differential vector. *F* serves as the differential weight or scaling parameter, selected from the range of 0 to 2. Moreover, 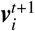 denotes the *i*th component of vector ***v*** at time *t* + 1.
3. Crossover: The crossover operator is governed by the parameter *CR* ∈ [0, 1], representing the probability parameter for the Crossover. For the entire population (*i* = 1, …, *n*), let *j*_*rand*_ = ⌊*rand*[0, 1) × *D*⌋ and for all dimensions (*j* = 1, …, *D*), update the *j*th component from any mutated vector ***v***_*i*_ with potential crossover operation as follows:

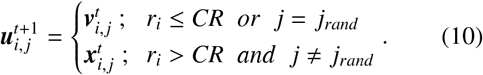

In this case, *r*_*i*_ is a random value ranging from 0 to 1. Notably, this crossover step is linked to DE with binomial crossover, which is akin to GA with a one-point crossover. DE with exponential crossover is also equivalent to GA with two-point crossover, but it is not employed in the proposed approach.
4. Selection: Across the entire population (*i* = 1, …, *n*), the solution ***x***_*i*_ is selected and updated through assessment of the trial vector 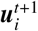 and optimization of the objective function *f* (.) as:

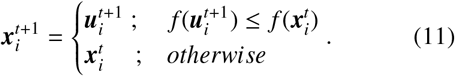

This four-step process is iterated across all generations.

## 3. Proposed Method

Machine learning techniques are widely recognized as a fundamental approach for motor imagery processing. In this MI-BCI study, we integrate classical signal processing with GSP techniques to finely construct and reduce the brain graph, selecting optimal features derived from the graph. This section describes the stages and procedures employed for preprocessing, classical signal processing, and GSP. Fig. 1 illustrates the block diagram of our proposed method, referred to as K-GLR-DE.

**Figure 1:**
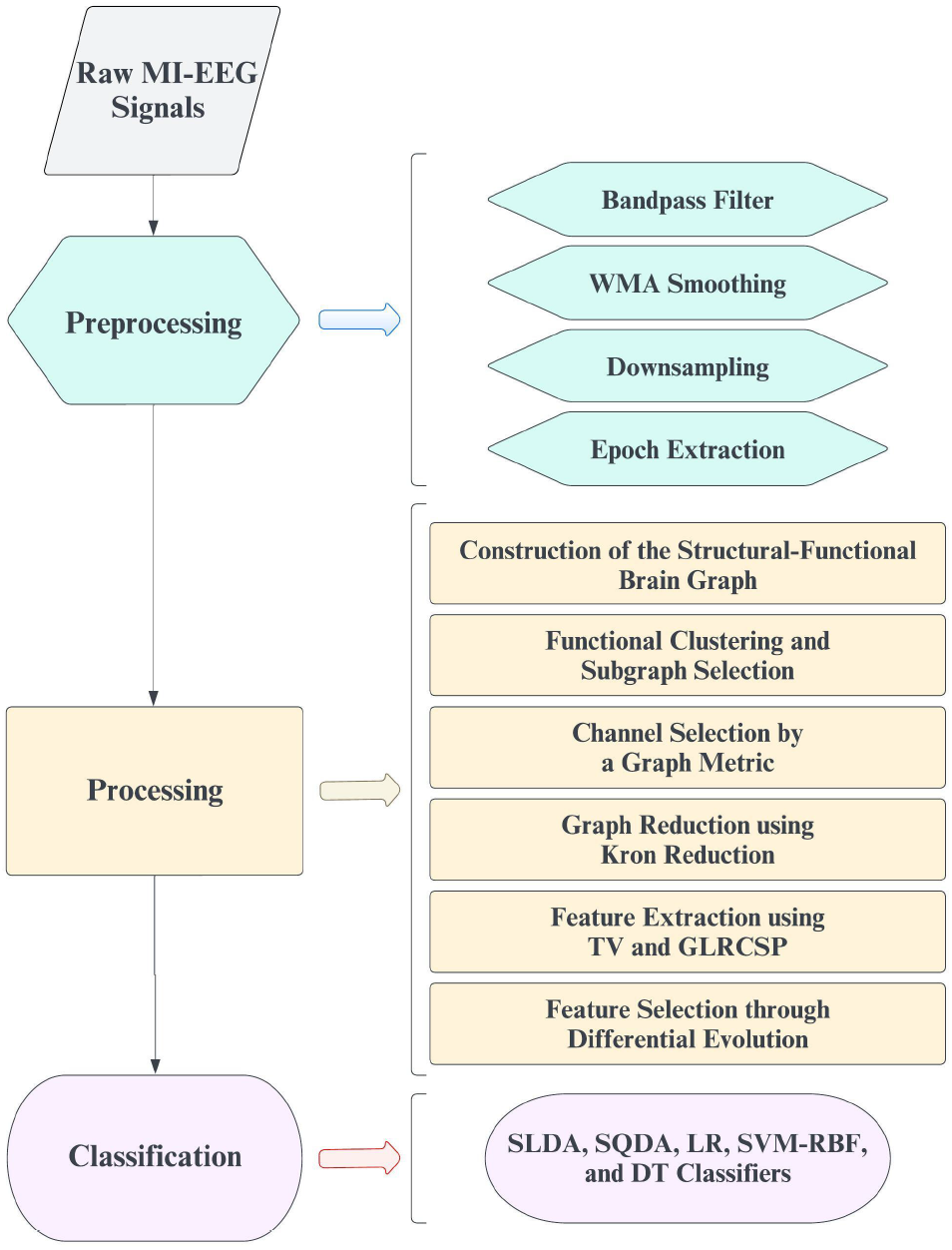
Block diagram of the proposed method, K-GLR-DE

### 3.1. Preprocessing

For better signal processing, we employ a fifth-order Butterworth band-pass filter within the range of 8-30 Hz, guided by previous studies [26, 54], to preserve the alpha (8-13 Hz) and beta (13-30 Hz) frequency bands. Reliable studies have demonstrated the significant importance of these two frequency bands in classifying MI-BCI tasks [54, 55]. As indicated by [26], we employ the weighted moving average (WMA) method to account for the subject’s response to each stimulus. Through the examination of various WMA functions in this application, we find that the square function yields lower reconstruction error and superior results compared to linear and binomial functions.

Numerous studies on BCI systems, including [24-26, 56], have employed downsampling as an effective preprocessing technique. A fixed downsampling factor of 10 is used, reducing the sampling frequency from 1000 Hz to 100 Hz. To mitigate the interference of visual evoked potential preceding the EEG signal, certain studies consider 0.5 s after stimulus application as a valuable window to examine the MI-EEG [30]. To extract MI activity, similar to [24-26], we analyze the time interval of 0.5 s to 4 s following the visual stimulus in each trial of the dataset.

### 3.2. Processing

In the BCI system, EEG signals undergo processing to extract the essential control commands. This paper employs classical signal processing and GSP as the primary approaches. The following steps and methods are explained: graph construction, graph reduction, feature extraction, feature selection, and classification.

#### 3.2.1. Construction of the Structural-Functional Brain Graph

As described in Section 2.3, the construction of graphs, including brain graphs, necessitates the identification of two fundamental components: vertices and weighted edges. Expanding on our previous study [28], this paper investigates the utilization of EEG signals on structural-functional brain graphs (SFG brain graphs). The vertices in our brain network correspond to the EEG electrodes, resulting in 118 vertices. The weights of brain graph edges are determined by measures of structural connectivity through geometric distance and functional connectivity through correlation, following a methodology akin to [28]. Within the SFG brain graphs, edge weights are determined as the product of two exponential expressions, representing the normalized squares of the Euclidean distance and the Pearson correlation. The weight matrix ***W***_SFG_ for *p* = 1, …, *N* and *q* = 1, …, *N*, is defined as:

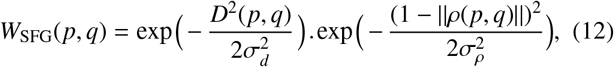

Where *D*(*p, q*) *< k* represents an element from the Euclidean distance matrix, and *ρ*(*p, q*) represents an element from the correlation matrix. In (12), *σ*_*d*_ and *σ*_*ρ*_ denote the normalization coefficients for the squared Euclidean distance and squared Pearson correlation between vertices, respectively. The value of *k* represents the threshold for the Euclidean distance between two vertices. If this distance falls below the threshold *k*, the weight of the edge connecting the two electrodes is calculated using (12); otherwise, the weight is set to 0. In this study, following the approach of [28], we determine the optimal normalizing coefficients *σ*_*d*_ and *σ*_*ρ*_ for each subject based on three criteria: the constant number (as per [24]), the variance of matrix elements, and the maximum of matrix elements. The threshold value *k* is set to 1 (according to the head radius), as stipulated by [24]. Consequently, employing (12), we compute the weighted adjacency matrix, which possesses dimensions of (118 × 118), for the SFG brain graphs. Subsequently, we acquire the graph Laplacian matrix, which has dimensions of (118 × 118), for use in the ensuing procedures. Graphs are extracted individually for each subject but collectively for the two types of MI activities.

#### 3.2.2. Functional Clustering and Subgraph Selection

To identify the active brain regions during MI tasks, we consider the specific physiology of the case and apply a functional clustering method to the EEG electrodes. Functional clustering is performed based on the Brodmann areas atlas, corresponding to 118 EEG channels in the current application. Additionally, we consider the somatosensory cortical areas responsible for the MI tasks to improve classification performance [57, 58]. Fig. 2 (a) displays the twenty-four Regions of Interest (ROIs) for various types of MI tasks. Fig. 2 (b) illustrates the functional clustering of 118 EEG electrodes based on 24 ROIs, which correspond to 12 bilateral ROIs encompassing the left and right hemispheres. The details of all 24 ROIs are provided in Table 2 [57, 58]. The number of electrodes considered in each subgraph is defined as follows:

**Table 2:**
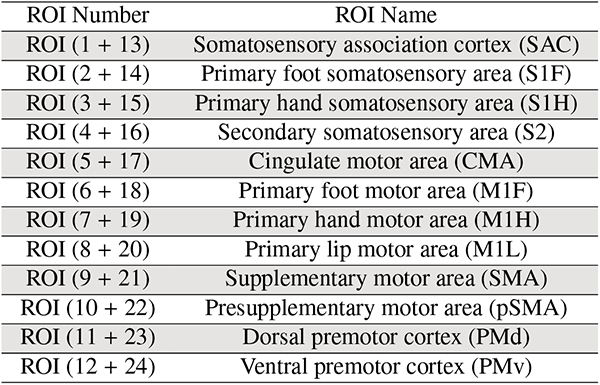
24 ROIs in MI-BCI.

**Figure 2:**
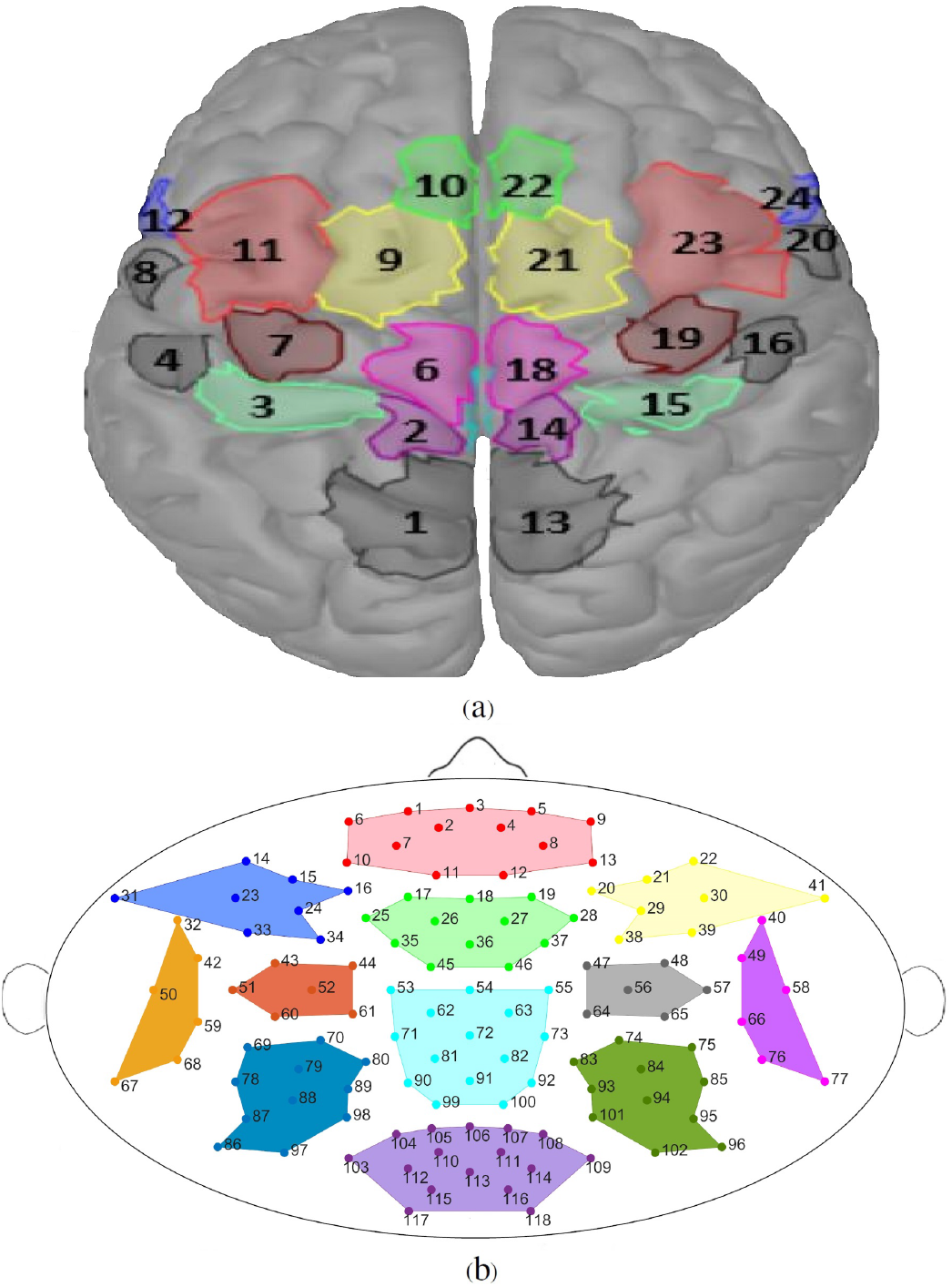
(a): 24 ROIs for MI tasks [58], (b): Functional clustering of 118 EEG electrodes based on specific ROIs for hand-foot MI tasks

**Figure 3:**
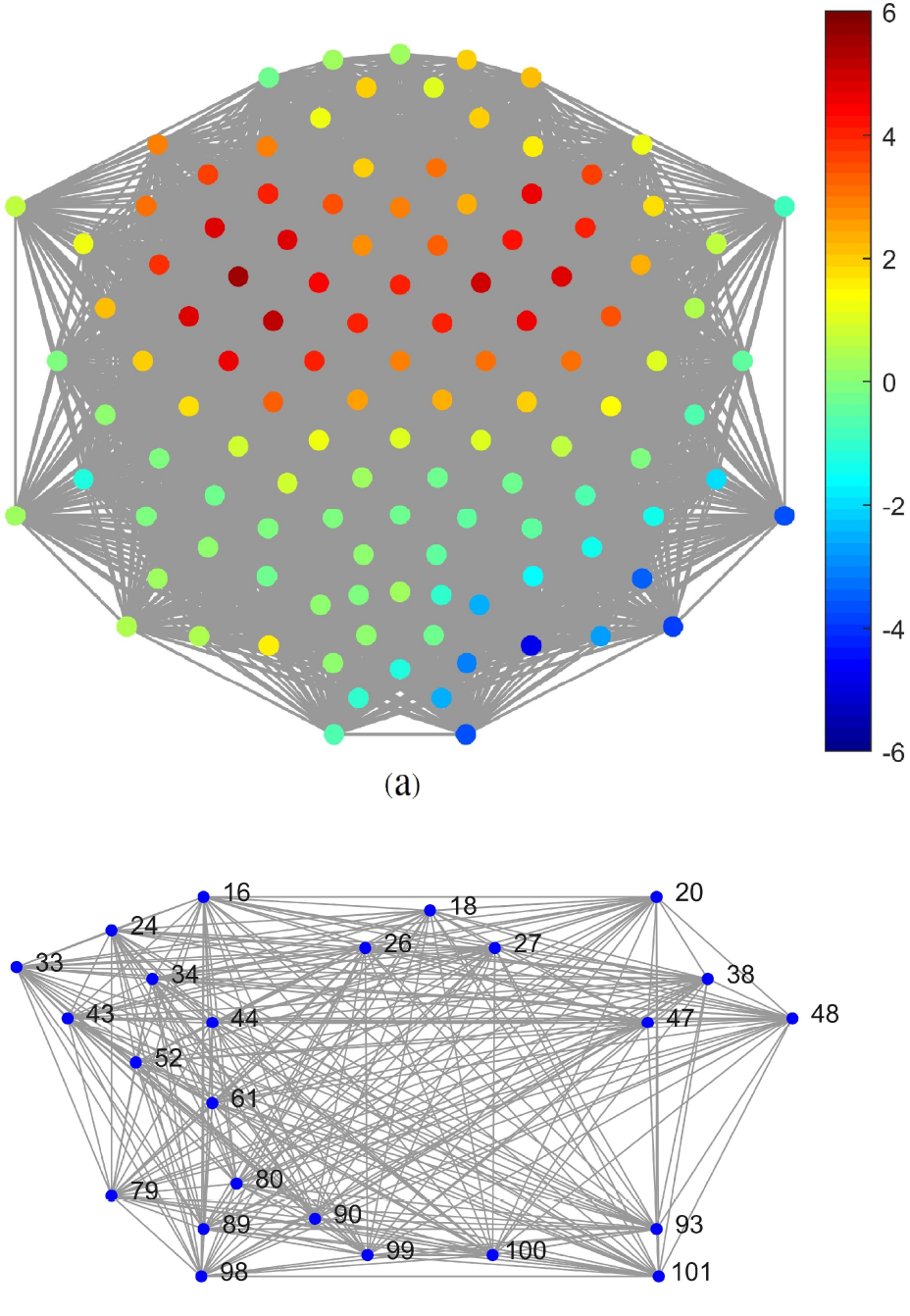
(a): Example of an EEG signal on the original SFG brain graph for subject aa in the vertex domain, (b): Example of the reduced SFG brain graph for subject aa (24 significant vertices) achieved through weighted degree and Kron reduction in the vertex domain.

- Cluster 1: [1-13],
- Cluster 2: [17-19, 25-28, 35-37, 45-46],
- Cluster 3: [14-16, 23-24, 31, 33-34],
- Cluster 4: [20-22, 29-30, 38-39, 41],
- Cluster 5: [32, 42, 50, 59, 67-68],
- Cluster 6: [40, 49, 58, 66, 76-77],
- Cluster 7: [43-44, 51-52, 60-61],
- Cluster 8: [47-48, 56-57, 64-65],
- Cluster 9: [53-55, 62-63, 71-73, 81-82, 90-92, 99-100],
- Cluster 10: [69-70, 78-80, 86-89, 97-98],
- Cluster 11: [74-75, 83-85, 93-96, 101-102],
- Cluster 12: [103-118].

The analysis was conducted within 18 specific regions, which encompassed the right hand and right foot tasks, including pairs such as {R2 + R14, R3 + R15, R5 + R17, R6 + R18, R7 + R19, R9 + R21, R10 + R22, R11 + R23, R12 + R24}. As depicted in Fig. 2, these 18 ROIs are associated with eight subgraphs, specifically {C2, C3, C4, C7, C8, C9, C10, C11}. Accordingly, we identify significant physiological areas in this case and conduct preliminary channel selection by prioritizing more important subgraphs, resulting in the selection of 77 out of 118 vertices.

#### 3.2.3. Channel Selection by a Graph Metric

Channel selection plays a crucial role in brain signal processing as it reduces computational complexity. Choosing the appropriate channels in EEG processing is analogous to selecting important vertices in the brain graph from the perspective of GSP. Centrality metrics are a set of criteria in network science that examines the relative importance of a vertex or an edge concerning the structure and function of the network. Therefore, centralities are influential criteria for selecting vertices. Weighted degree, closeness centrality, betweenness centrality, eigenvector centrality, vulnerability centrality, and PageRank centrality are among the most prominent and widely used centrality metrics in brain networks and social networks [59]. In this step, we employ a widely adopted and straightforward metric known as the weighted degree to select 24 significant vertices from the eight related brain subgraphs, which are then utilized in the Kron reduction process to simplify the graph. According to neurophysiological characteristics [48, 58, 60], MI-EEG of the right hand and right foot occurs more frequently in the brain’s left hemisphere than in the right hemisphere. Based on our experimental observations, we have found that assigning greater weight to the left hemisphere of the brain in this context produces superior outcomes. Instead of selecting the three top vertices from each of the eight subgraphs, we choose four superior vertices from three left brain subgraphs, three superior vertices from two middle brain subgraphs, and two superior vertices from three right brain subgraphs, resulting in a total of 24 superior vertices in the brain graph.

#### 3.2.4. Graph Reduction using Kron Reduction

This study addresses the challenge of dimension reduction in large EEG data through Kron reduction. To reduce the vertex dimension in the brain graph, we employ a combination of selecting significant vertices based on degree centrality and utilizing Kron reduction. By utilizing degree centrality and Kron reduction based on (2) and (3), we select 24 superior vertices from eight related subgraphs. Consequently, we reduce the number of vertices in eight subgraphs from 77 to 24 and the size of the Laplacian matrix from (77 × 77) to (24 × 24) in the vertex domain. This approach enables meaningful sampling in the vertex domain by merging the selection of the top 24 vertices from the initial 77 vertices and applying Kron reduction to the Laplacian matrix. Fig.3 (a) illustrates an example of an EEG signal on the SFG brain graph in the vertex domain for subject aa, while Fig.3 (b) demonstrates the reduced SFG brain graph achieved through weighted degree and Kron reduction for subject aa.

#### 3.2.5. Feature Extraction using TV and GLRCSP

In this study, we used *TV*_𝒢_(***x***) as GSP-based feature vectors. After Kron reduction, ***W*** has dimensions of (24 × 24) in (1). We apply the collected data separately for the 24 vertices of eight subgraphs as described in (1). Consequently, the feature vectors for each subgraph and each trial will have dimensions of (350 × 1).

Inspired by findings from [48], we employ GLRCSP method to regularize spatial filters in our study. CSP-based feature extraction was conducted by inputting EEG data from eight subgraphs into GLRCSP with *m* spatial filters. Similar to the methodology outlined in [48], we meticulously optimized hyperparameters to achieve maximum 10-fold cross-validation accuracy on the training dataset, leveraging LDA as the classifier. Values for *β* and *γ* were chosen from the set {0, 0.1, 0.2, …, 0.9}.

Overall, feature extraction contained two distinctive categories: 1) Total variation derived from eight subgraphs, yielding dimensions of 8 × 350 per trial, and 2) The logarithm of the variance of each GLCRSP output to the logarithm of the total variance of all GLRCSP outputs. This resulted in dimensions of *m* × 1 per trial, where *m* ∈ {2, 4} spatial filters were applied. These categories encapsulate a set of graph-based features in the vertex domain and statistical features. Consequently, feature vectors were consolidated from these two distinct feature categories. Subsequently, feature selection was executed through the implementation of DE.

#### 3.2.6. Feature Selection through Differential Evolution

As elucidated in Section 3.2.5, the number of features per trial is too large for the number of trials. However, this count proves excessive for robust classification within our dataset comprising 280 trials, potentially triggering the curse of dimensionality. In response, we leverage an approach aimed at optimizing feature selection while effectively reducing feature dimensions. This involves the application of an approximate optimization technique known as differential evolution.

Similar to [49, 50], the optimization objective function *f* (.) equates to the percentage of classification accuracy for training data. The training process of this research has duly taken this function into account. A noteworthy advantage of the DE algorithm lies in its utilization of differential vectors, capturing the disparity between responses and incorporating valuable insights from excluded individuals in the population. In this study, our objective function is focused on classification accuracy. The parameters employed for the DE algorithm are presented in Table 3. Feature selection via the DE algorithm encompasses two key components: first, the construction of a feature subset through DE, and second, the utilization of a classification algorithm to assess the validity of the chosen subset. Ultimately, the DE algorithm’s output yields a selection of the top 10 superior features, which are then employed for classification.

**Table 3:**
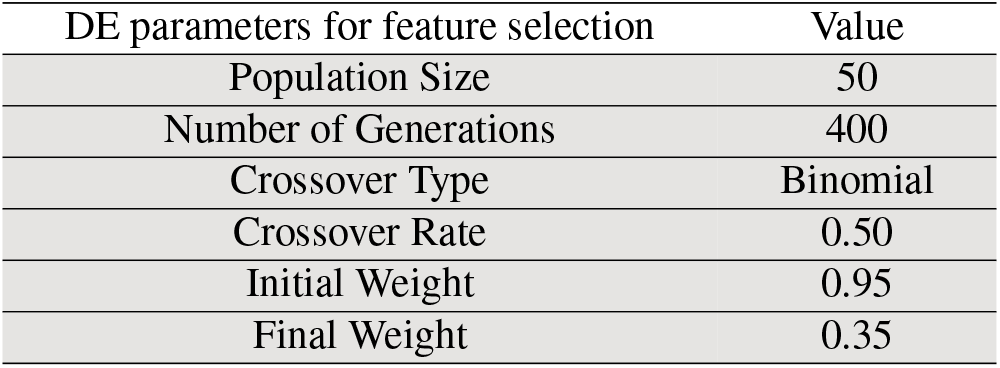
Parameters used in DE for selecting the optimal set of 10-features.

### 3.3. Classification

Following the feature selection process, we proceed to the classification stage. In MI-BCI processing, our primary objective is accurately assign samples to their classes. To evaluate the performance of our proposed approach (K-GLR-DE), the features identified by the DE algorithm are utilized as input for five classifiers: shrinkage LDA (SLDA), shrinkage QDA (SQDA), logistic regression (LR), SVM with radial basis function (SVM-RBF), and decision tree (DT). Validation is carried out using a 10-fold cross-validation method.

The SLDA classifier, an extension of LDA, incorporates regularization through the shrinkage of class-related covariance matrices [61]. This regularization is crucial because covariance matrices estimated from small datasets often exhibit extreme eigenvalues, leading to inaccurate covariance estimates. This challenge is mitigated by applying regularization to covariance matrices ∑, resulting in 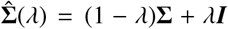 where ***I*** represents the identity matrix and *λ* is the regularization parameter. Notably, analytical solutions exist for the automatic selection of the optimal *λ* value. In ERP-based BCI [61] and oscillatory activity BCI [62], the resulting SLDA classifier has demonstrated superiority over the conventional LDA classifier. Similarly, the SQDA classifier, a variant of QDA, is also subject to regularization for enhanced performance.

## 4. Results and Discussion

This section comprehensively presents the quantitative and intuitive analysis of outcomes across diverse scenarios in our proposed methodology, facilitated through tables and charts. These scenarios encompass: A) Evaluating the impact of Kron reduction within the contexts of limited, small, and conventional training data, along with the combined influence of Kron reduction and an increased number of GLRCSP spatial filters; B) Examining the implications of Kron reduction in the context of BCI Competition III-Dataset IVa training trials; and C) Investigating the combined effect of Kron reduction and differential evolution. We delve into a detailed discussion and analysis of the accuracy achieved by all five classifiers (Section 3.3) under varying conditions, providing insights into the separability assessment of the two-class motor imagery tasks (right hand and right foot).

### 4.1. Impact of Kron Reduction on Limited, Small, and Conventional Training Trials

Given the significance of data size in classification, we thoroughly analyze the effects of Kron reduction across three distinct training data size modes, two spatial filter modes, and a comprehensive trend analysis within this section.

#### 4.1.1. Comparative Analysis of GLR-DE and GDR-BCI [26] Results

Table 4 presents the mean outcomes for GLR-DE and the mean results of GDR-BCI’s best fold across all five subjects, employing LDA, QDA, and LR classifiers in six distinct modes. Upon reviewing Table 4, it is evident that the average accuracy of LDA, QDA, and LR has shown improvement in this study compared to the average accuracy of the best fold for these three classifiers in [26] across all six modes. Additionally, the standard deviation of GLR-DE accuracy is lower than that of GDR-BCI accuracy, signifying the enhanced stability of our recommended approach over the GDR-BCI method [26].

**Table 4:** Comparative analysis of GLR-DE and GDR-BCI [26] results across six different modes (Number of spatial filters-Number of training trials)

| Scenarios | LDA | LDA-[26] | QDA | QDA-[26] | LR | LR-[26] |
| --- | --- | --- | --- | --- | --- | --- |
| (2-60) | 72.23 | 64.73 | 71.35 | 63.36 | 73.02 | 64.52 |
| (4-60) | 78.94 | 74.36 | 79.09 | 70.91 | 79.64 | 73.91 |
| (2-100) | 82.86 | 76.42 | 81.74 | 78.00 | 82.92 | 76.89 |
| (4-100) | 85.43 | 78.22 | 85.28 | 78.00 | 86.77 | 78.91 |
| (2-200) | 87.11 | 80.50 | 87.07 | 80.25 | 87.77 | 80.75 |
| (4-200) | 88.22 | 80.25 | 88.50 | 80.75 | 89.13 | 81.00 |

#### 4.1.2. Limited (60) Training Trials with 2 and 4 GLRCSP Spatial Filters

Table 5-(a), (b) present results for (2-60) GLR-DE and (2-60) K-GLR-DE, while Fig. 4 illustrates the impact of Kron reduction in (2-60). Similarly, Table 6-(a), (b) depict findings for (4-60) GLR-DE and (4-60) K-GLR-DE, highlighting Kron reduction efficacy in the (4-60) mode through Fig. 5. Applying the K-GLR-DE approach to 2-GLRCSP and limited (60) training trials yielded average accuracy improvements of 2.01%, 2.19%, 2.17%, 1.93%, and 2.16%, for the SLDA, SQDA, LR, SVM-RBF, and DT classifiers compared to GLR-DE, respectively. Similarly, in the case of 4-GLRCSP and limited (60) training trials, applying the K-GLR-DE approach led to average accuracy enhancements of 3.21%, 3.22%, 3.12%, 3.28%, and 3.10% for the same classifiers compared GLR-DE, respectively. As a result, the Kron reduction applied to the GLR-DE approach increased the average accuracy of all five classifiers in both (2-60) and (4-60) scenarios.

**Table 5:** Classification accuracy (Mean±Std) with two spatial filters and 60 training trials employing GLR-DE and K-GLR-DE.

| (a)<br>(2-60) GLR-DE |  |  |  |  |  |
| --- | --- | --- | --- | --- | --- |
| Subjects | SLDA | SQDA | LR | SVM-RBF | DT |
| aa | 68.26±1.40 | 68.35±0.98 | 67.43±1.80 | 72.62±2.04 | 71.04±0.88 |
| al | 83.29±0.78 | 82.00±0.74 | 82.45±1.46 | 85.74±1.48 | 84.88±1.22 |
| av | 67.93±3.54 | 67.84±1.83 | 67.45±4.15 | 71.90±3.02 | 70.02±2.44 |
| aw | 71.85±2.04 | 71.09±0.98 | 72.31±2.04 | 76.31±2.13 | 74.08±1.62 |
| ay | 75.74±1.98 | 76.04±1.28 | 75.48±2.14 | 79.61±2.48 | 78.48±2.10 |
| Mean | 73.41±1.87 | 73.06±1.16 | 73.02±2.32 | 77.24±2.23 | 75.70±1.65 |

| (b)<br>(2-60) K-GLR-DE |  |  |  |  |  |
| --- | --- | --- | --- | --- | --- |
| Subjects | SLDA | SQDA | LR | SVM-RBF | DT |
| aa | 71.00±1.76 | 70.75±1.57 | 70.41±1.69 | 74.99±1.88 | 73.14±1.79 |
| al | 84.28±0.71 | 84.00±0.76 | 84.07±1.37 | 86.92±1.26 | 86.07±0.59 |
| av | 70.25±3.25 | 70.39±1.82 | 69.91±3.28 | 73.72±2.58 | 73.15±2.03 |
| aw | 74.87±2.33 | 74.08±0.88 | 74.89±0.92 | 79.54±2.04 | 77.37±1.39 |
| ay | 76.69±1.56 | 77.04±0.93 | 76.69±2.01 | 80.69±2.36 | 79.56±1.09 |
| Mean | 75.42±1.92 | 75.25±1.19 | 75.19±1.85 | 79.17±2.02 | 77.86±1.38 |

**Table 6:** Classification accuracy (Mean±Std) with four spatial filters and 60 training trials employing GLR-DE and K-GLR-DE.

| (a) |  |  |  |  |  |
| --- | --- | --- | --- | --- | --- |
| (4-60) GLR-DE |  |  |  |  |  |
| Subjects | SLDA | SQDA | LR | SVM-RBF | DT |
| aa | 74.81±3.55 | 76.76±3.24 | 73.85±3.66 | 77.87±2.65 | 77.07±2.23 |
| al | 86.99±0.70 | 87.23±1.33 | 85.37±1.54 | 89.79±2.01 | 89.34±2.13 |
| av | 74.38±4.24 | 75.71±4.18 | 73.83±4.41 | 76.80±3.60 | 76.03±2.84 |
| aw | 81.40±2.51 | 83.31±2.49 | 79.98±2.67 | 84.61±2.84 | 83.77±2.72 |
| ay | 86.40±0.97 | 86.16±2.17 | 85.18±1.94 | 88.69±2.25 | 87.59±1.64 |
| Mean | 80.80±2.39 | 81.83±2.68 | 79.64±2.84 | 83.55±2.67 | 82.75±2.31 |

| (b) |  |  |  |  |  |
| --- | --- | --- | --- | --- | --- |
| (4-60) K-GLR-DE |  |  |  |  |  |
| Subjects | SLDA | SQDA | LR | SVM-RBF | DT |
| aa | 78.50±2.87 | 80.75±3.10 | 77.47±3.12 | 81.21±2.76 | 80.41±2.90 |
| al | 89.88±1.00 | 90.43±1.55 | 89.22±1.65 | 92.73±1.74 | 92.32±1.91 |
| av | 77.66±3.55 | 78.43±3.50 | 75.70±3.94 | 80.50±3.26 | 78.64±3.07 |
| aw | 83.95±2.04 | 85.67±2.55 | 82.51±2.22 | 86.98±2.52 | 86.09±2.57 |
| ay | 90.04±1.39 | 89.98±2.14 | 88.90±2.04 | 92.73±2.52 | 91.80±2.28 |
| Mean | 84.01±2.17 | 85.05±2.57 | 82.76±2.59 | 86.83±2.56 | 85.85±2.55 |

**Figure 4:**
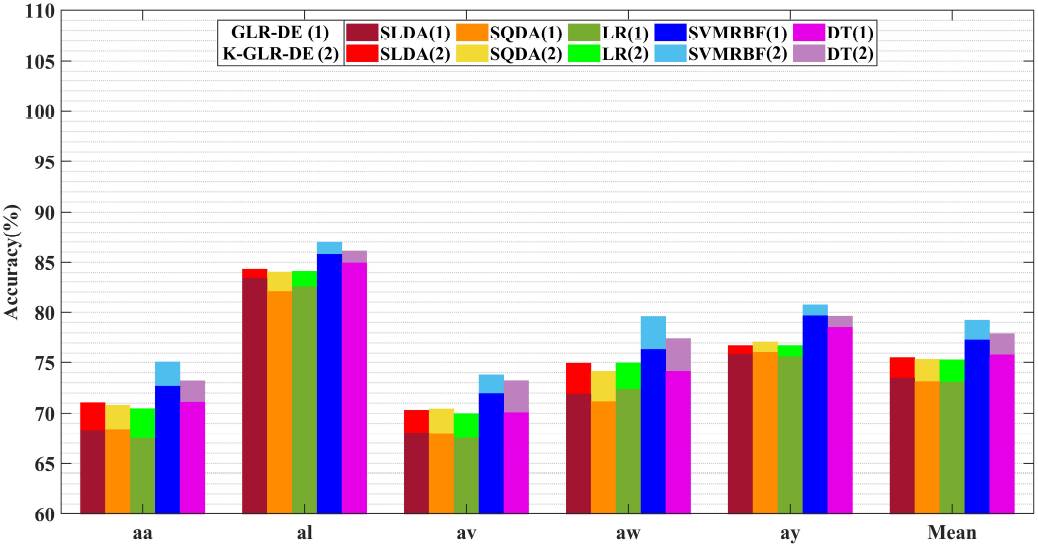
Effect of Kron reduction on classification accuracy in (2-60) GLR-DE and (2-60) K-GLR-DE

**Figure 5:**
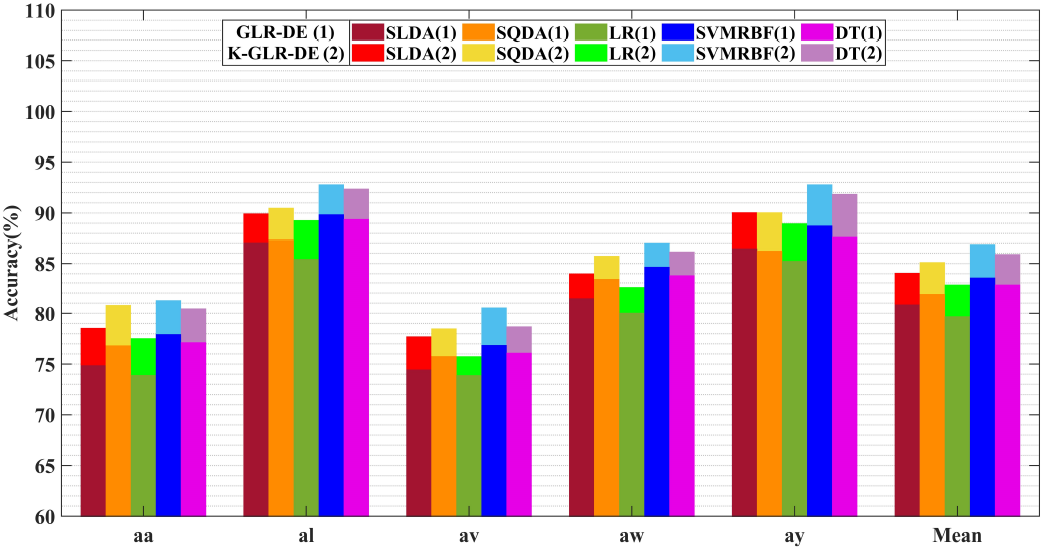
Effect of Kron reduction on classification accuracy in (4-60) GLR-DE and (4-60) K-GLR-DE

#### 4.1.3. Small (100) Training Trials with 2 and 4 GLRCSP Spatial Filters

Table 7-(a), (b) present the outcomes of (2-100) GLR-DE and (2-100) K-GLR-DE, respectively, while Fig. 6 illustrates the impact of Kron reduction within the (2-100) context. Similarly, Table 8-(a), (b) detail the findings of (4-100) GLR-DE and (4-100) K-GLR-DE, respectively, showcasing the effectiveness of Kron reduction in the (4-100) mode as illustrated in Fig. 7. In the (2-100) scenario, the K-GLR-DE approach yielded mean improvements of 2.86%, 3.27%, 3.48%, 3.91%, and 3.53% in the performance of SLDA, SQDA, LR, SVM-RBF, and DT classifiers, respectively, compared to the GLR-DE approach. Correspondingly, in the (4-100) scenario, the K-GLR-DE approach led to average enhancements of 3.40%, 2.99%, 1.23%, 2.09%, and 2.07% in the same classifiers, compared to the GLR-DE approach. Consequently, the application of the K-GLR-DE approach led to heightened mean accuracy across all five classifications for both the (2-100) and (4-100) cases, surpassing the performance of the GLR-DE approach.

**Table 7:** Classification accuracy (Mean±Std) with two spatial filters and 100 training trials employing GLR-DE and K-GLR-DE.

| (a) |  |  |  |  |  |
| --- | --- | --- | --- | --- | --- |
| (2-100) GLR-DE |  |  |  |  |  |
| Subjects | SLDA | SQDA | LR | SVM-RBF | DT |
| aa | 85.98±1.30 | 84.09±1.76 | 83.78±1.50 | 88.10±2.17 | 86.64±1.40 |
| al | 90.09±1.24 | 87.13±1.58 | 89.70±1.27 | 92.11±0.74 | 91.30±1.10 |
| av | 73.08±1.97 | 74.22±1.87 | 71.84±1.69 | 77.55±2.13 | 76.51±1.64 |
| aw | 86.95±1.17 | 86.24±1.34 | 86.20±1.52 | 88.94±1.59 | 88.39±1.36 |
| ay | 84.14±1.12 | 83.82±0.95 | 83.07±0.85 | 86.57±1.38 | 85.45±0.76 |
| Mean | 84.05±1.36 | 83.10±1.50 | 82.92±1.37 | 86.65±1.60 | 85.66±1.25 |

| (b) |  |  |  |  |  |
| --- | --- | --- | --- | --- | --- |
| (2-100) K-GLR-DE |  |  |  |  |  |
| Subjects | SLDA | SQDA | LR | SVM-RBF | DT |
| aa | 88.50±1.63 | 87.21±2.14 | 87.12±1.85 | 90.93±2.45 | 89.49±1.63 |
| al | 91.45±1.52 | 90.32±1.94 | 91.36±1.59 | 94.72±0.96 | 92.78±1.33 |
| av | 75.70±2.37 | 75.75±2.26 | 74.42±2.06 | 79.92±2.41 | 78.92±1.90 |
| aw | 90.56±1.67 | 90.48±1.67 | 90.74±1.49 | 94.98±1.62 | 93.76±1.02 |
| ay | 88.32±1.48 | 88.07±1.24 | 88.37±1.18 | 92.23±1.85 | 90.99±1.60 |
| Mean | 86.91±1.73 | 86.37±1.85 | 86.40±1.63 | 90.56±1.86 | 89.19±1.50 |

**Table 8:** Classification accuracy (Mean±Std) with four spatial filters and 100 training trials employing GLR-DE and K-GLR-DE.

| (a) |  |  |  |  |  |
| --- | --- | --- | --- | --- | --- |
| (4-100) GLR-DE |  |  |  |  |  |
| Subjects | SLDA | SQDA | LR | SVM-RBF | DT |
| aa | 85.11±2.64 | 85.17±1.96 | 85.56±1.86 | 88.20±2.43 | 87.63±1.66 |
| al | 93.28±1.37 | 93.78±0.52 | 93.29±0.90 | 96.31±1.80 | 95.76±1.42 |
| av | 78.02±3.48 | 79.05±2.98 | 79.14±2.74 | 81.95±2.60 | 81.67±2.54 |
| aw | 88.78±2.87 | 88.57±1.29 | 87.37±0.98 | 91.12±2.23 | 89.60±1.58 |
| ay | 87.54±1.83 | 87.30±1.90 | 87.32±2.12 | 91.41±1.84 | 90.62±1.80 |
| Mean | 86.55±2.44 | 86.37±1.73 | 86.77±1.72 | 89.80±2.18 | 89.06±1.80 |

| (b) |  |  |  |  |  |
| --- | --- | --- | --- | --- | --- |
| (4-100) K-GLR-DE |  |  |  |  |  |
| Subjects | SLDA | SQDA | LR | SVM-RBF | DT |
| aa | 89.05±2.83 | 87.77±2.16 | 86.56±2.36 | 90.64±2.21 | 89.61±2.16 |
| al | 96.06±1.57 | 96.04±0.71 | 94.46±1.11 | 98.28±1.26 | 97.17±1.43 |
| av | 80.81±3.18 | 79.63±3.18 | 79.58±2.94 | 83.07±3.43 | 82.17±3.12 |
| aw | 91.97±2.04 | 91.10±1.79 | 90.37±1.18 | 93.37±2.29 | 92.99±2.04 |
| ay | 91.33±3.07 | 92.24±1.90 | 89.05±2.32 | 94.08±2.60 | 93.72±2.34 |
| Mean | 89.95±2.54 | 89.36±1.95 | 88.00±1.98 | 91.89±2.36 | 91.13±2.22 |

**Figure 6:**
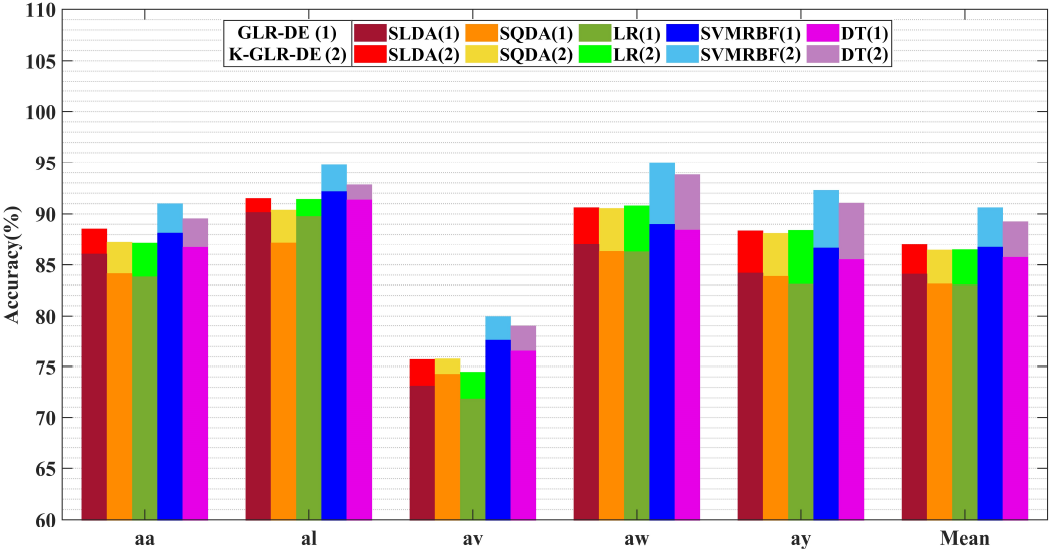
Effect of Kron reduction on classification accuracy in (2-100) GLR-DE and (2-100) K-GLR-DE

**Figure 7:**
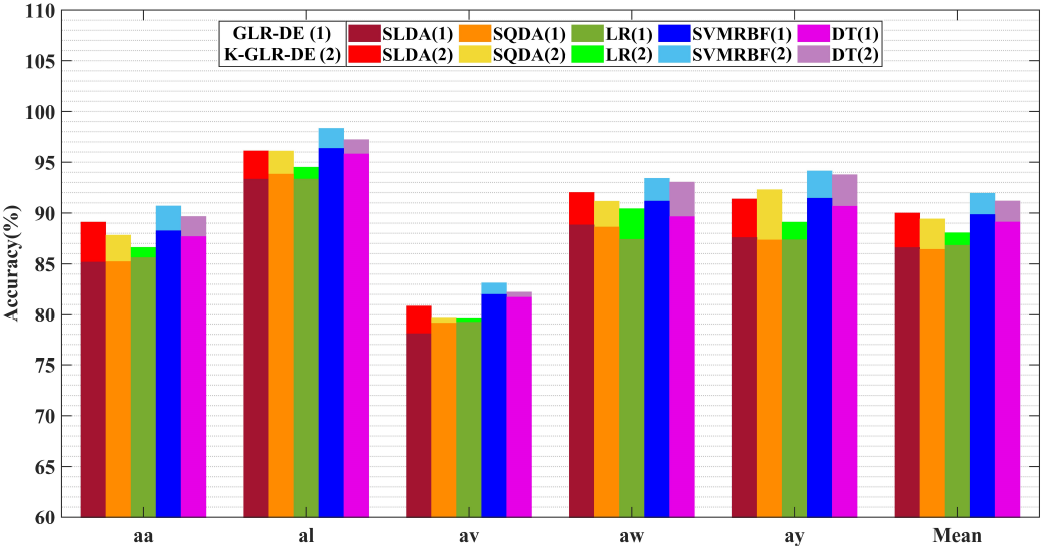
Effect of Kron reduction on classification accuracy in (4-100) GLR-DE and (4-100) K-GLR-DE

#### 4.1.4. Conventional (200) Training Trials with 2 and 4 GLRCSP Spatial Filters

Table 9-(a), (b) provide the outcomes for (2-200) GLR-DE and (2-200) K-GLR-DE, respectively, while Fig. 8 illustrates the impact of Kron reduction in this context. Similarly, Table 10-(a), (b) present the results of (4-200) GLR-DE and (4-200) K-GLR-DE, respectively, showcasing the effectiveness of Kron reduction in the (4-200) mode, as demonstrated in Fig. 9. Within the context of 2-GLRCSP mode and conventional training data (200), the mean accuracy of all subjects in SLDA, SQDA, LR, SVM-RBF, and DT classifiers using the K-GLR-DE approach has exhibited increments of 1.99%, 1.86%, 1.51%, 1.27%, and 0.99%, respectively, when compared to the GLR-DE approach. Similarly, in the scenario involving 4-GLRCSP mode and conventional training data (200), the K-GLR-DE approach has led to average enhancements of 2.03%, 2.06%, 2.77%, 2.39%, and 2.54% across the same classifiers, as opposed to the GLR-DE approach. Consequently, the application of Kron reduction has resulted in improved mean accuracy across all five classifiers in both the (2-200) and (4-200) cases.

**Table 9:** Classification accuracy (Mean±Std) with two spatial filters and 200 training trials employing GLR-DE and K-GLR-DE.

| (a) |  |  |  |  |  |
| --- | --- | --- | --- | --- | --- |
| (2-200) GLR-DE |  |  |  |  |  |
| Subjects | SLDA | SQDA | LR | SVM-RBF | DT |
| aa | 89.37±1.42 | 87.96±1.52 | 87.67±1.44 | 91.31±2.07 | 90.68±1.75 |
| al | 93.98±1.38 | 93.05±1.16 | 92.49±0.94 | 96.19±1.46 | 95.82±1.68 |
| av | 78.96±1.54 | 78.81±1.90 | 75.91±1.48 | 82.26±2.42 | 81.47±2.06 |
| aw | 92.53±0.63 | 92.92±0.63 | 92.73±0.91 | 96.19±1.35 | 95.54±1.24 |
| ay | 90.50±1.46 | 90.99±1.40 | 90.03±1.22 | 93.60±1.97 | 92.81±1.61 |
| Mean | 89.07±1.29 | 88.75±1.32 | 87.77±1.20 | 91.91±1.85 | 91.26±1.67 |

| (b) |  |  |  |  |  |
| --- | --- | --- | --- | --- | --- |
| (2-200) K-GLR-DE |  |  |  |  |  |
| Subjects | SLDA | SQDA | LR | SVM-RBF | DT |
| aa | 91.64±0.98 | 89.93±0.98 | 88.57±1.00 | 92.84±1.80 | 91.95±1.21 |
| al | 95.57±0.90 | 94.61±0.74 | 94.65±0.68 | 97.35±1.30 | 96.44±0.92 |
| av | 80.61±1.21 | 80.46±1.56 | 77.31±1.44 | 84.00±2.13 | 83.17±1.33 |
| aw | 94.74±0.48 | 94.18±0.38 | 94.01±0.52 | 97.03±1.32 | 96.44±0.94 |
| ay | 92.74±1.12 | 93.85±1.37 | 91.86±0.96 | 94.69±1.76 | 93.25±1.30 |
| Mean | 91.06±0.94 | 90.61±1.01 | 89.28±0.92 | 93.18±1.66 | 92.25±1.14 |

**Table 10:** Classification accuracy (Mean±Std) with four spatial filters and 200 training trials employing GLR-DE and K-GLR-DE.

| (a) |  |  |  |  |  |
| --- | --- | --- | --- | --- | --- |
| (4-200) GLR-DE |  |  |  |  |  |
| Subjects | SLDA | SQDA | LR | SVM-RBF | DT |
| aa | 89.40±1.03 | 89.34±0.82 | 88.09±1.21 | 92.44±1.13 | 91.45±1.03 |
| al | 96.55±1.00 | 96.23±0.60 | 95.76±1.10 | 98.66±0.49 | 97.35±0.79 |
| av | 82.40±1.44 | 82.11±1.26 | 81.04±1.62 | 85.45±1.25 | 84.51±2.21 |
| aw | 93.13±0.80 | 94.20±0.71 | 91.39±1.46 | 96.68±1.10 | 95.23±1.58 |
| ay | 91.77±1.38 | 94.13±1.02 | 89.38±1.36 | 95.07±0.96 | 94.29±1.01 |
| Mean | 90.68±1.13 | 91.18±0.88 | 89.13±1.35 | 93.66±0.99 | 92.57±1.32 |

| (b) |  |  |  |  |  |
| --- | --- | --- | --- | --- | --- |
| (4-200) K-GLR-DE |  |  |  |  |  |
| Subjects | SLDA | SQDA | LR | SVM-RBF | DT |
| aa | 90.72±1.75 | 91.08±1.95 | 90.27±2.13 | 95.93±1.44 | 94.87±1.22 |
| al | 96.80±1.84 | 97.35±1.51 | 97.50±1.23 | 99.55±0.45 | 98.55±0.80 |
| av | 85.11±2.26 | 85.03±2.33 | 83.71±2.08 | 88.73±1.70 | 89.46±1.48 |
| aw | 96.22±1.62 | 96.77±1.67 | 94.78±1.72 | 99.10±0.89 | 96.12±1.04 |
| ay | 94.71±2.04 | 96.04±1.53 | 93.26±1.98 | 96.95±1.14 | 96.55±1.01 |
| Mean | 92.71±1.90 | 93.24±1.80 | 91.90±1.75 | 96.05±1.22 | 95.11±1.11 |

**Figure 8:**
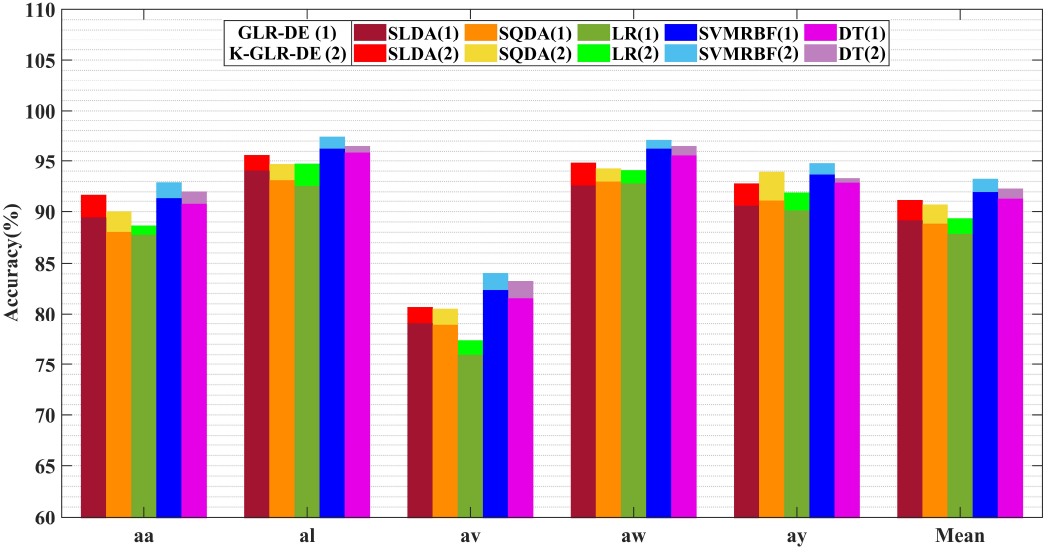
Effect of Kron reduction on classification accuracy in (2-200) GLR-DE and (2-200) K-GLR-DE

**Figure 9:**
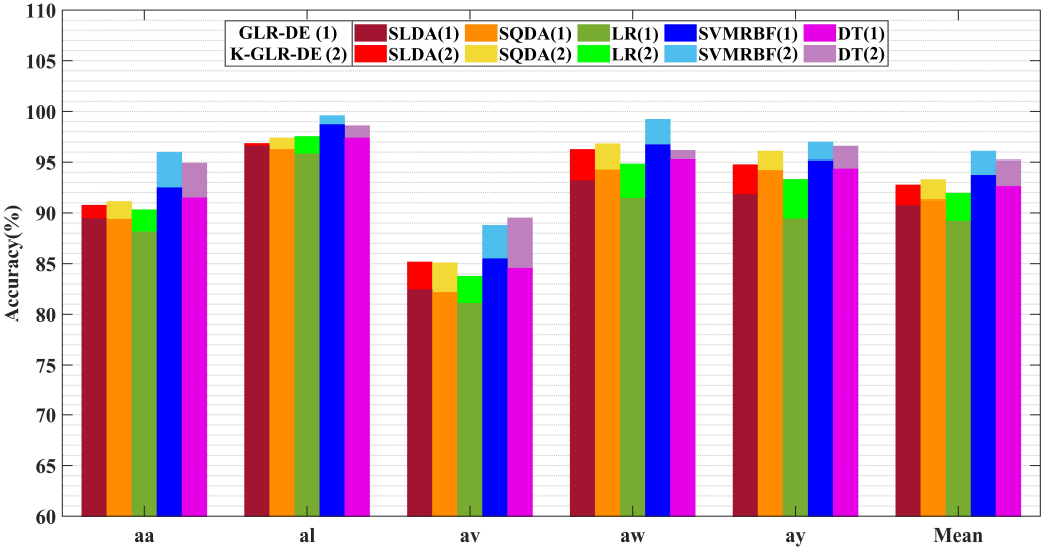
Effect of Kron reduction on classification accuracy in (4-200) GLR-DE and (4-200) K-GLR-DE

Notably, the investigation conducted in this study reveals that the average accuracy of SLDA and SQDA adaptive classifiers consistently surpasses that of the fixed LDA and QDA cases across all modes. This includes scenarios with limited, small, and conventional training datasets, and configurations involving two and four spatial filters.

#### 4.1.5. Trends in Training Trials and Spatial Filters

Figures 10 (a) and 10 (b) visually represent the fluctuations in the average accuracy of the best classifier in this study, namely SVM-RBF, concerning the number of training trials. The variations are shown for both GLR-DE and K-GLR-DE approaches within the 2-GLRCSP mode. Across the 2-GLRCSP mode, both GLR-DE and K-GLR-DE approaches exhibit an ascending trend in SVM-RBF accuracy as the number of trials increases from 60 to 100 and subsequently to 200. In Fig. 10 (a), the SVM-RBF accuracy for aa and aw demonstrates a marked rise from 60 to 100 trials, followed by a slight increase from 100 to 200 trials. Conversely, for al, av, and ay, the augmentation in average SVM-RBF accuracy from 60 to 100 and 100 to 200 trials remains relatively constant. In Fig. 10 (b), SVM-RBF accuracy for aa, aw, and ay experiences a steep ascent from 60 to 100 trials and a moderate rise from 100 to 200 trials. However, the increase in SVM-RBF accuracy for al and av from 60 to 100 and 100 to 200 trials is more consistent. Overall, the trend indicates a favorable impact of increased training trials on SVM-RBF accuracy within the 2-GLRCSP mode for both GLR-DE and K-GLR-DE approaches.

**Figure 10:**
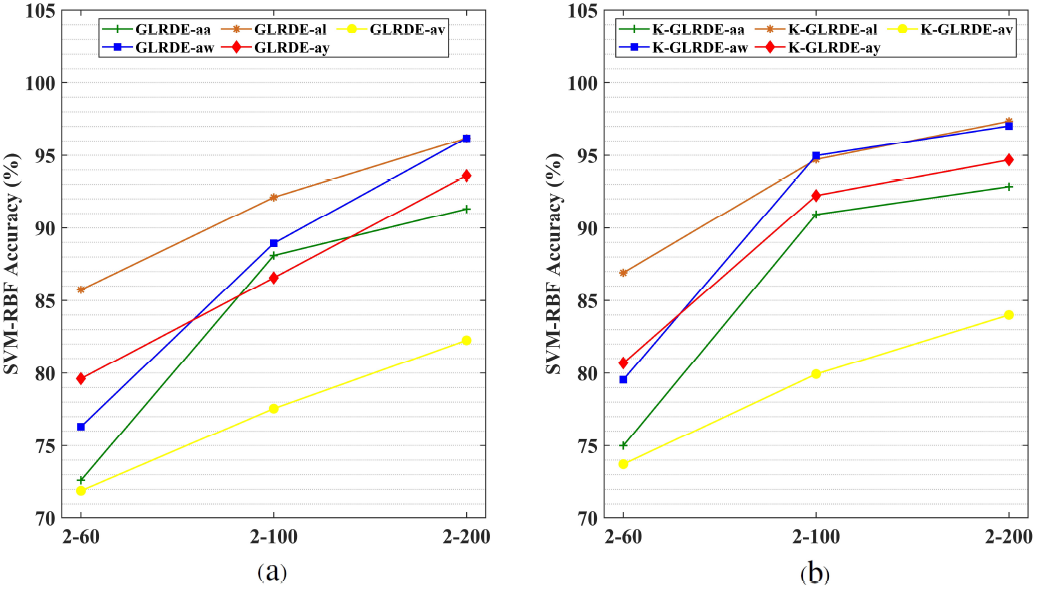
Trend of (60-100-200) training trials with two spatial filters in GLR-DE and K-GLR-DE approaches

Figures 11 (a) and 11 (b) present the variations in average SVM-RBF accuracy as the number of training trials changes, comparing GLR-DE and K-GLR-DE approaches in the context of 4-GLRCSP. Moreover, in the 4-GLRCSP mode, both the GLR-DE and K-GLR-DE methods exhibit an increase in average SVM-RBF accuracy by augmenting the trial count from 60 to 100 and subsequently to 200. In Fig. 11 (a), the average SVM-RBF accuracy for aa, al, av, and aw demonstrates a noteworthy rise from 60 to 100 trials and a moderate rise from 100 to 200 trials. However, for ay, the increase in average SVM-RBF accuracy from 60 to 100 trials exceeds that from 100 to 200 trials. In Fig. 11 (b), the average SVM-RBF accuracy for aa and al has substantially increased from 60 to 100 trials, followed by a gradual increase from 100 to 200 trials. On the contrary, the slope of the chart for aw remains relatively constant between 60 to 100 and 100 to 200 trials. Notably, the augmentation in average SVM-RBF accuracy for av and ay from 60 to 100 trials surpasses that from 100 to 200 trials.

**Figure 11:**
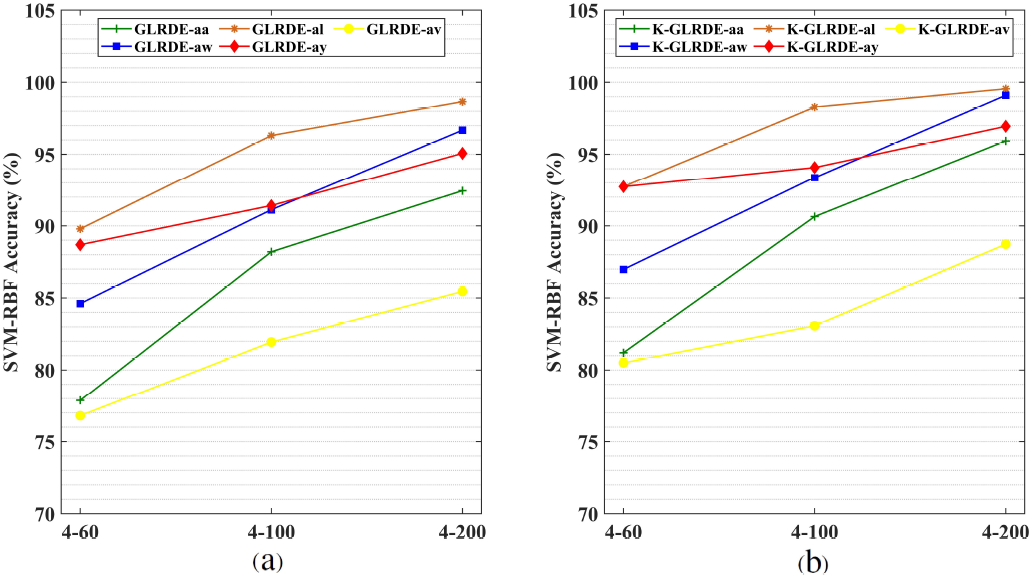
Trend of (60-100-200) training trials with four spatial filters in GLR-DE and K-GLR-DE approaches

### 4.2. Impact of Kron Reduction on Training Trials of BCI Competition III-Dataset IVa

To provide a comparative analysis with the outcomes from the BCI Competition III-dataset IVa, we adopt similar experimental conditions. In the 4-GLRCSP mode, aligned with the number of training trials in BCI Competition-III, the GLR-DE approach yields average accuracies of 87.14, 88.21, 86.02, 90.20, and 89.56 for LDA, QDA, LR, SVM-RBF, and DT classifiers, respectively (Table 11, Fig. 12). Similarly, under the same settings, the K-GLR-DE approach produces average accuracies of 90.39, 90.34, 88.80, 93.02, and 91.81 for the aforementioned classifiers (Table 12, Fig. 13). This demonstrates that incorporating Kron reduction leads to enhanced average accuracy across all classifiers. Moreover, the average performance of the K-GLR-DE method with SVM-RBF is noteworthy. In comparison with the top three performing groups in the BCI competition III-dataset IVa, who achieved respective average accuracies of 94.17, 85.12, and 83.45 [29], the competitiveness of the K-GLR-DE results becomes evident. Additional insights can be found in Table 12 and Fig. 13.

**Table 11:** Classification accuracy (Mean±Std) for 4-GLR-DE approach using training trials of BCI competition III-dataset IVa.

| 4-GLR-DE (BCI competition III-dataset IVa) |  |  |  |  |  |
| --- | --- | --- | --- | --- | --- |
| Subjects | SLDA | SQDA | LR | SVM-RBF | DT |
| aa (112) | 87.94±1.50 | 90.42±1.58 | 86.28±1.65 | 90.84±1.99 | 89.59±1.71 |
| al (56) | 94.83±0.99 | 96.68±0.95 | 94.82±1.29 | 98.54±1.16 | 98.33±1.21 |
| av (224) | 76.22±1.92 | 76.70±1.90 | 74.84±1.81 | 78.56±2.35 | 78.08±2.23 |
| aw (196) | 91.38±1.25 | 91.91±1.22 | 89.77±1.47 | 94.03±1.62 | 93.70±1.62 |
| ay (252) | 85.31±1.66 | 85.32±1.38 | 84.39±1.57 | 89.03±1.95 | 88.10±1.88 |
| Mean | 87.14±1.46 | 88.21±1.41 | 86.02±1.56 | 90.20±1.81 | 89.56±1.73 |

**Table 12:** Classification accuracy (Mean±Std) for 4-K-GLR-DE approach using training trials of BCI competition III-dataset IVa.

| 4-K-GLR-DE (BCI competition III-dataset IVa) |  |  |  |  |  |
| --- | --- | --- | --- | --- | --- |
| Subjects | SLDA | SQDA | LR | SVM-RBF | DT |
| aa (112) | 88.89±0.84 | 91.37±0.96 | 87.25±0.85 | 91.78±1.19 | 90.13±1.15 |
| al (56) | 98.35±0.53 | 97.47±0.58 | 97.88±0.65 | 99.95±0.05 | 98.84±0.88 |
| av (224) | 78.93±1.04 | 79.23±1.11 | 77.53±1.16 | 81.24±1.36 | 80.31±1.45 |
| aw (196) | 94.91±0.67 | 93.69±0.90 | 92.33±0.88 | 98.29±0.96 | 97.58±1.07 |
| ay (252) | 90.85±0.69 | 89.92±0.92 | 89.00±0.69 | 93.83±0.98 | 92.17±1.04 |
| Mean | 90.39±0.75 | 90.34±0.89 | 88.80±0.93 | 93.02±0.91 | 91.81±1.12 |

**Figure 12:**
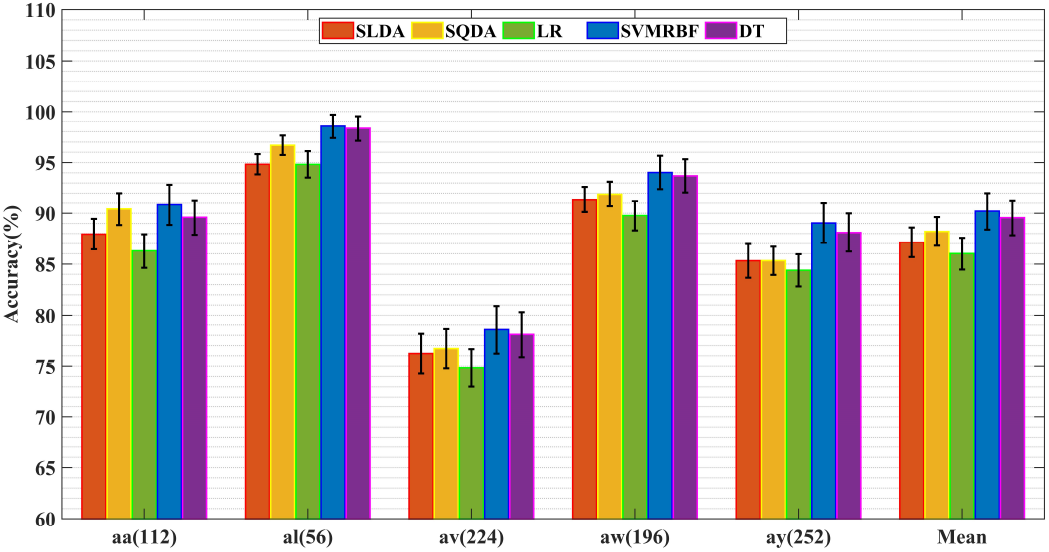
Effect of Kron reduction on classification accuracy (Mean±Std) for BCI competition III-dataset IVa training trials in 4-GLR-DE approach

**Figure 13:**
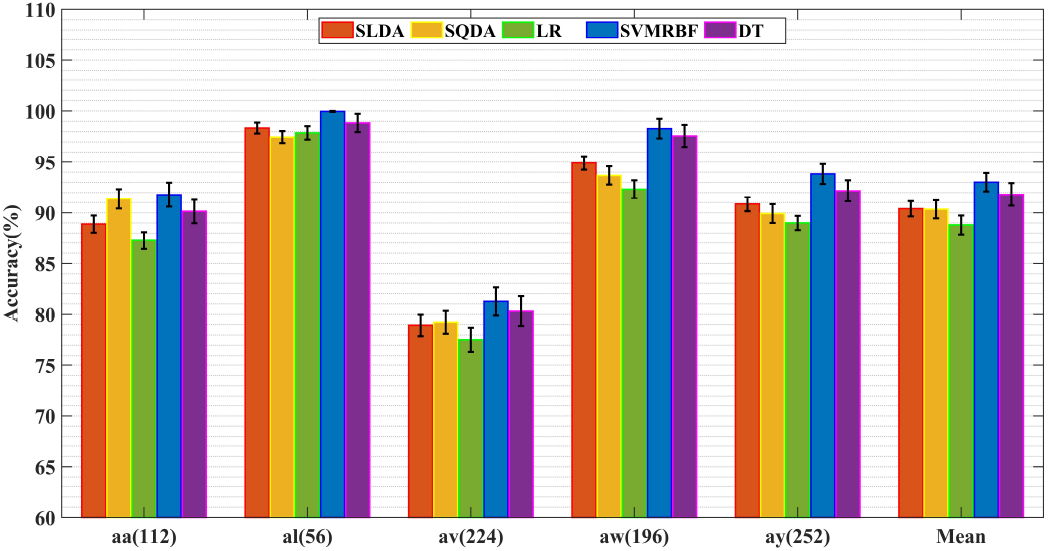
Effect of Kron reduction on classification accuracy (Mean±Std) for 4-K-GLR-DE approach using training trials of BCI competition III-dataset IVa

When analyzing result dispersion, it becomes evident from Fig. 12 and Fig. 13 that the standard deviation of av results surpasses that of other subjects, while for al, it is lower compared to the other subjects, signifying more stability in the al data results. This trend is also apparent in Tables 5-10.

Further insights can be drawn from Tables 11 and 12, which reveal that, despite having a similar number of test trials as BCI Competition-III, the classification outcomes for al and aw outperform those of other subjects. This might be attributed to the data recording process, wherein al and aw underwent two sessions of task 1 (target-correlated eye movement) and two sessions of task 2 (target-uncorrelated eye movement), in contrast to the other subjects who underwent one session of task 1 and three sessions of task 2. This pattern aligns with findings in previous studies as well.

### 4.3. Synergistic Impact of Kron Reduction and DE Algorithm

In this section, we explore the combined effect of Kron reduction and the DE algorithm on K-GLR-DE, aiming to draw comparisons with two earlier studies [28, 49]. The outcomes of the K-GLR-DE approach, evaluated through a 10-fold cross-validation across all 280 trials encompassing different individuals and classifiers, are documented in Table 13. Notably, Table 13 highlights that the K-GLR-DE approach coupled with SVM-RBF classification yields the highest average accuracy across all subjects. Moreover, the analysis in Table 13 reveals a noteworthy observation: the number of selected features for all subjects in [49] exceeds the ten features selected in this study and our previous work [28]. Additionally, our investigation notes that within the K-GLR-DE context, the SVM and DT classifiers exhibit superior accuracy compared to the SLDA, SQDA, and LR classifiers.

**Table 13:** Classification accuracy (Mean±Std) in 4-K-GLR-DE approach with 10-fold CV compared to KG-DE [28] and CSP-DE [49].

| 4-K-GLR-DE (10-fold CV) |  |  |  |  |  |  |  |
| --- | --- | --- | --- | --- | --- | --- | --- |
| Subjects | SLDA | SQDA | LR | SVM-RBF | DT | SVM-RBF[28] | SVM[49] |
| aa | 90.90±0.84 | 91.20±0.96 | 90.40±0.85 | 96.10±1.20 | 93.80±1.15 | 94.20±1.36 | 95.80±2.40 |
| al | 96.90±0.53 | 97.60±0.58 | 97.60±0.65 | 99.70±0.19 | 98.70±0.88 | 99.30±0.61 | 98.80±0.79 |
| av | 85.20±1.04 | 84.80±1.11 | 83.30±1.16 | 89.90±1.36 | 89.50±1.45 | 88.50±2.43 | 89.80±3.36 |
| aw | 95.60±0.67 | 97.20±0.90 | 96.40±0.88 | 99.40±0.43 | 96.40±1.07 | 98.60±0.94 | 99.20±0.72 |
| ay | 95.90±0.69 | 97.10±0.92 | 93.50±0.69 | 97.20±0.98 | 96.90±1.04 | 96.90±1.03 | 96.50±1.11 |
| Mean | 92.90±0.75 | 93.58±0.89 | 92.24±0.93 | 96.46±0.83 | 95.06±1.12 | 95.50±1.27 | 96.02±1.68 |

The average performance of SVM-RBF using the K-GLR-DE approach demonstrated an increase of 0.44 compared to the CSP-DE method [49] under the same 10-fold cross-validation scenario. Moreover, when compared to the KG-DE approach [28], the increase was even more substantial at 0.96. To provide a more intuitive and comprehensive difference of the outcomes, statistical summaries of classification accuracy for various participants are visually depicted as box plots in both Fig. 14 and Fig. 15.

**Figure 14:**
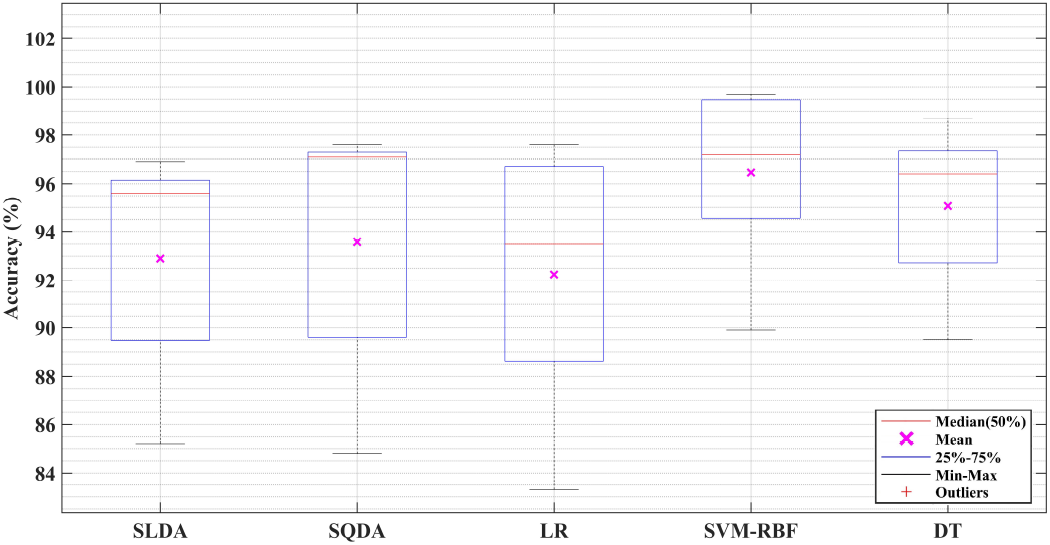
Accuracy of classifiers in 4-K-GLR-DE approach with 10-fold CV

**Figure 15:**
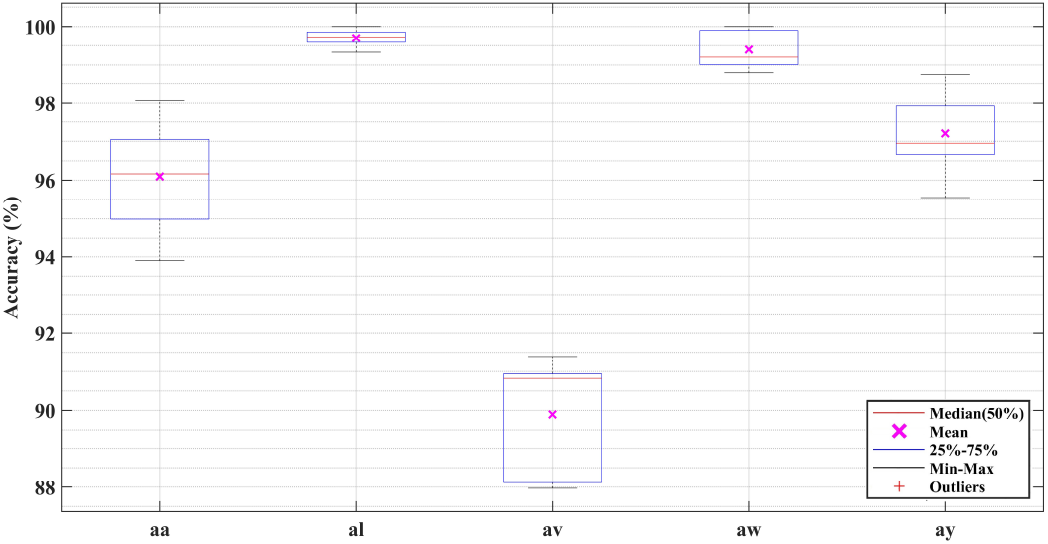
Accuracy of subjects in 4-K-GLR-DE approach with 10-fold CV

Figures 14 and 15 pertain to the K-GLR-DE method with a 10-fold cross-validation. In Fig. 14, the box plots represent an overview of the average results attained by the 4-K-GLR-DE method across all subjects and various classifiers. In Fig. 15, the box plots depict the average results achieved by the 4-K-GLR-DE method for the best classifier (SVM-RBF) and different subjects.

Both box plots (Figures 14 and 15) present essential statistical measures—mean, median, first and third quartiles, maximum, and minimum. These values are meticulously provided to facilitate an intuitive and comprehensive analysis of statistical variations in classifier performance, individual accuracy through the top classifier, and the effectiveness of the K-GLR-DE method itself. Notably, a trend consistent with previous studies on this dataset emerges: subject al demonstrates the highest classifier performance, while subject av exhibits the lowest. SVM-RBF and DT within the K-GLR-DE framework stand out for their notable performance.

### 4.4. Computational Efficiency in K-GLR-DE Approach for BCI System

In the K-GLR-DE methodology tailored for the BCI system, optimization parameters are computed during distinct phases of the training process for individual subjects. These calculated optimal coefficients and parameters are then utilized in subsequent test operations, ensuring a streamlined and rapid process. This efficiency contributes to shorter test execution times while achieving consistently high levels of accuracy.

We recommend using simple yet effective classifiers, including SLDA, SQDA, and DT, as they tend to yield promising outcomes. Both our research findings and [63] support this recommendation. Notably, SLDA and SQDA stand out due to their absence of hyperparameters, which enhances their user-friendliness. It is important to acknowledge that in cases involving extensive training data, the linear nature of LDA might lead to suboptimal results. In contrast, DT—a non-linear classifier—performs admirably across diverse training set sizes. In terms of computational complexity, DT, akin to SLDA and SQDA, emerges as an efficient and swift approach that has proven successful in online applications.

In the most time-consuming scenario in our study, each classification label for a 3.5-second test trial is determined approximately 60 milliseconds after trial completion. We conducted the experiments using MATLAB 2018b on a laptop featuring an Intel Core i7-6700 2.6 GHz CPU and 16 GB RAM. Given the preceding considerations, the execution time of the K-GLR-DE test process falls within a range conducive for practical utilization in online BCI systems.

## 5. Conclusions

This paper introduces the K-GLR-DE approach, designed to achieve dimension reduction and classification of EEG in MI-BCI. The approach encompasses several key components: subgraph selection based on physiological ROIs, Kron reduction for dimensionality reduction of the graph Laplacian matrix, total variation and GLRCSP for extracting features, and the DE optimization algorithm for selecting and reducing feature dimensions.

Kron reduction, aided by selecting the crucial vertices, considers the Laplacian matrix data from all brain graph vertices. The Kron reduction efficiently streamlines Laplacian graph information by incorporating selected and deleted vertices. GLRCSP, a generalized method of CSP, adeptly captures the CSP-based features of selected channels by regulating parameters *β* and *γ*. The DE algorithm, a meta-heuristic evolutionary technique, emerges as an excellent choice for feature selection, offering superior convergence, approximating global optima, ease of use, and success across practical problems.

The combination of GSP-based total variation features and usual CSP-based statistical features in MI, achieved through DE-enhanced feature selection, yields improved classification accuracy. The addition of GSP-based features to CSP-based features yields superior performance in MI classification compared to using only CSP-based features.

In terms of classification outcomes, adaptive classification methods and spatial filters demonstrate advantages for their efficiency and robustness with smaller datasets. Significantly, SLDA and SQDA outperform their traditional counterparts, LDA and QDA, within the scope of this study. Moreover, DT demonstrates its merit as a reliable choice, while SVM emerges as a widely used and remarkably performing classifier in BCI systems.

The results highlight the improved performance resulting from Kron reduction coupled with physiology-graph-based channel selection for MI classification, particularly in limited, small, and conventional training trials. The K-GLR-DE approach, in conjunction with SVM-RBF and DT classifiers, achieves accuracy levels comparable to the top performers in BCI competition III-dataset IVa. Employing four spatial filters and SVM-RBF classification, the proposed approach achieves an average accuracy of 93.02 ± 0.91. Notably, SVM-RBF emerges as the best classifier within the K-GLR-DE framework, boasting an average accuracy of 96.46 ± 0.83 in the 4-K-GLR-DE setup. This signifies an enhancement of 0.44 over the CSP-DE method [49] and a substantial improvement of 0.96 compared to the KG-DE method in our previous study [28].

